# DNA metabarcoding provides insights into the diverse diet of a dominant suspension feeder, the giant plumose anemone *Metridium farcimen*

**DOI:** 10.1101/2020.12.07.407288

**Authors:** Christopher D. Wells, Gustav Paulay, Bryan N. Nguyen, Matthieu Leray

## Abstract

Benthic suspension feeders have significant impacts on plankton communities by depleting plankton or modifying composition of the plankton through prey selectivity. Quantifying diets of planktivorous animals can be difficult because plankton are frequently microscopic, may lack diagnostic characters, and are digested at variable rates. With DNA metabarcoding, the identification of gut contents has become faster and more accurate, and the technique allows for higher taxonomic resolution-n while also identifying rare and highly degraded items that would otherwise not be detected. We used DNA metabarcoding to examine the diet of the giant plumose anemone *Metridium farcimen,* a large, abundant, competitively-dominant anemone on subtidal rock surfaces and floating docks in the northeast Pacific Ocean. Gut contents of 12 individuals were compared to 80- and 330-μm filtered plankton samples collected one hour prior between 0.02 and 1.5 km from the anemones. The objectives of this study were to determine if *M. farcimen* has a selective diet and compare our findings with traditional gut content analyses. *Metridium farcimen* captured a wider range of prey than previously suspected and metabarcoding found many more taxa than traditional sampling techniques. Gut contents were less diverse than 80-μm filtered plankton samples, but more diverse than 330-μm filtered plankton samples. The diet of the anemones was 52% arthropods with a surprisingly high relative abundance of an ant (10%) that has mating flights in August when this study was conducted. The gut contents of *M. farcimen* likely include all prey that it can detect and that cannot escape. There were no overrepresented taxa in the gut contents compared to the plankton but there were underrepresented taxa. This study highlights the usefulness of the metabarcoding method in identifying prey within the gut of planktivorous animals and the significant terrestrial input into marine food webs.

## INTRODUCTION

Benthic suspension feeders can have major impacts on the structure of littoral food webs (Gili & Coma, 1998; Kimmerer, Gartside, & Orsi, 1994; Petersen, 2004; Sebens & Koehl, 1984; Whitten, Marin Jarrin, & McNaught, 2018; Young & Gotelli, 1988). Dense populations of suspension feeders can filter the immediately overlying water volume several times a day (Davies, Stuart, & de Villiers, 1989; Jørgensen, 1980; Petersen, 2004; Petersen & Riisgård, 1992; Riisgård, 1991; Vedel, Andersen, & Riisgård, 1994) until depleting it from resources (Riisgård, Jürgensen, & Clausen, 1996; Riisgård, Poulsen, & Larsen, 1996; Vedel, 1998).

Both passive and active benthic suspension feeders rely on water flow to bring food particles to their vicinity. Yet, all prey do not have equal probability of being captured. Prey use mechanisms such as escape behaviors, morphological defenses, and toxicity to avoid predation (Browman, Kruse, & O’Brien, 1989; Dodson, 1974; Engström, Viherluoto, & Viitasalo, 2001; Safi, Hewitt, & Talman, 2007; Suchman & Sullivan, 1998; Viitasalo, Flinkman, & Viherluoto, 2001; Viitasalo, Kiørboe, Flinkman, Pedersen, & Visser, 1998). In addition, numerous suspension feeders are highly selective for certain types of prey. Many bivalves, for example, preferentially select plankton based on size, shape, and motility (Cucci et al., 1985; Defossez & Hawkins, 1997; Safi et al., 2007; Shumway, Cucci, Newell, & Yentsch, 1985). Prey selectivity can result from the inability of predators to capture certain prey, the preferential capture or consumption of palatable and energetically valuable species, or active rejection of prey. Studies on the dietary selectivity of benthic suspension feeders are key to our understanding of the effects of predators on their ecosystem and of the role of dietary niche partitioning for species coexistence (Costello & Colin, 2002; Leray et al., 2019; Suchman & Sullivan, 1998).

Unfortunately, quantifying the diet of planktivorous animals can be difficult because plankton are frequently microscopic, often lack diagnostic characters, are digested quickly and at variable rates, and resources for identification are poorly developed for most regions (Fancett, 1988; Larson, 1991; Purcell, 1977; Sebens & Koehl, 1984; Zamer, 1986). Because of this, marine plankton are difficult to identify in gut contents past the class or order level when using traditional visual identification techniques (e.g., Fancett, 1988; Purcell, 1977; Sebens & Koehl, 1984).

With the advent of high throughput sequencing and powerful molecular techniques such as DNA metabarcoding (Taberlet, Coissac, Pompanon, Brochmann, & Willerslev, 2012), identification of specimens within community samples, such as in plankton and gut contents, can be rapid, accurate, and relatively cheap (Aylagas, Borja, & Rodríguez-Ezpeleta, 2014; Brandon-Mong et al., 2015; Nielsen, Clare, Hayden, Brett, & Kratina, 2018). DNA metabarcoding has been used to successfully identify taxa within gut contents of fishes (Albaina, Aguirre, Abad, Santos, & Estonba, 2016; Harms-Tuohy, Schizas, & Appeldoorn, 2016; Leray, Meyer, & Mills, 2015; Leray, Yang, et al., 2013), to evaluate biodiversity of insects (Brandon-Mong et al., 2015; Ji et al., 2013; Yu et al., 2012), and to identify the presence of rare taxa using environmental DNA (Deiner et al., 2017; Evans et al., 2016; Valentini et al., 2016). Metabarcoding can be used to analyze diets to reach a higher taxonomic resolution while also identifying rare and highly degraded items that would otherwise not be detected (Nielsen et al., 2018).

We used DNA metabarcoding to examine the diet of the giant plumose anemone *Metridium farcimen* (Brandt, 1835, Fig. 1). *Metridium farcimen* (Cnidaria, Anthozoa, Actiniaria) is a large, abundant sea anemone on subtidal rock surfaces and floating docks in the northeast Pacific Ocean (Fautin, Bucklin, & Hand, 1989; Fautin & Hand, 2000; Hand, 1955; Kozloff, 1973; Ricketts, Calvin, & Hedgpeth, 1968) that feeds on small zooplankton (Koehl, 1977a; Purcell, 1977; Sebens, 1981; Shick, 1991). This anemone, which can extend over a meter into the water column (Fautin et al., 1989), is well-adapted for high-flow environments (Koehl, 1977a, 1977b, 1977c) and is a competitive dominant species on rocky subtidal ledge communities (Nelson & Craig, 2011; Wells, 2019; Wells & Sebens, 2017). Gut contents of *M. farcimen* in the San Juan Archipelago, WA, USA were compared to concurrent, nearby plankton samples to determine if *M. farcimen* has a selective diet and to compare findings from traditional gut content analyses (i.e., suction and dissection) with metabarcoding analysis.

**Fig. 1.**
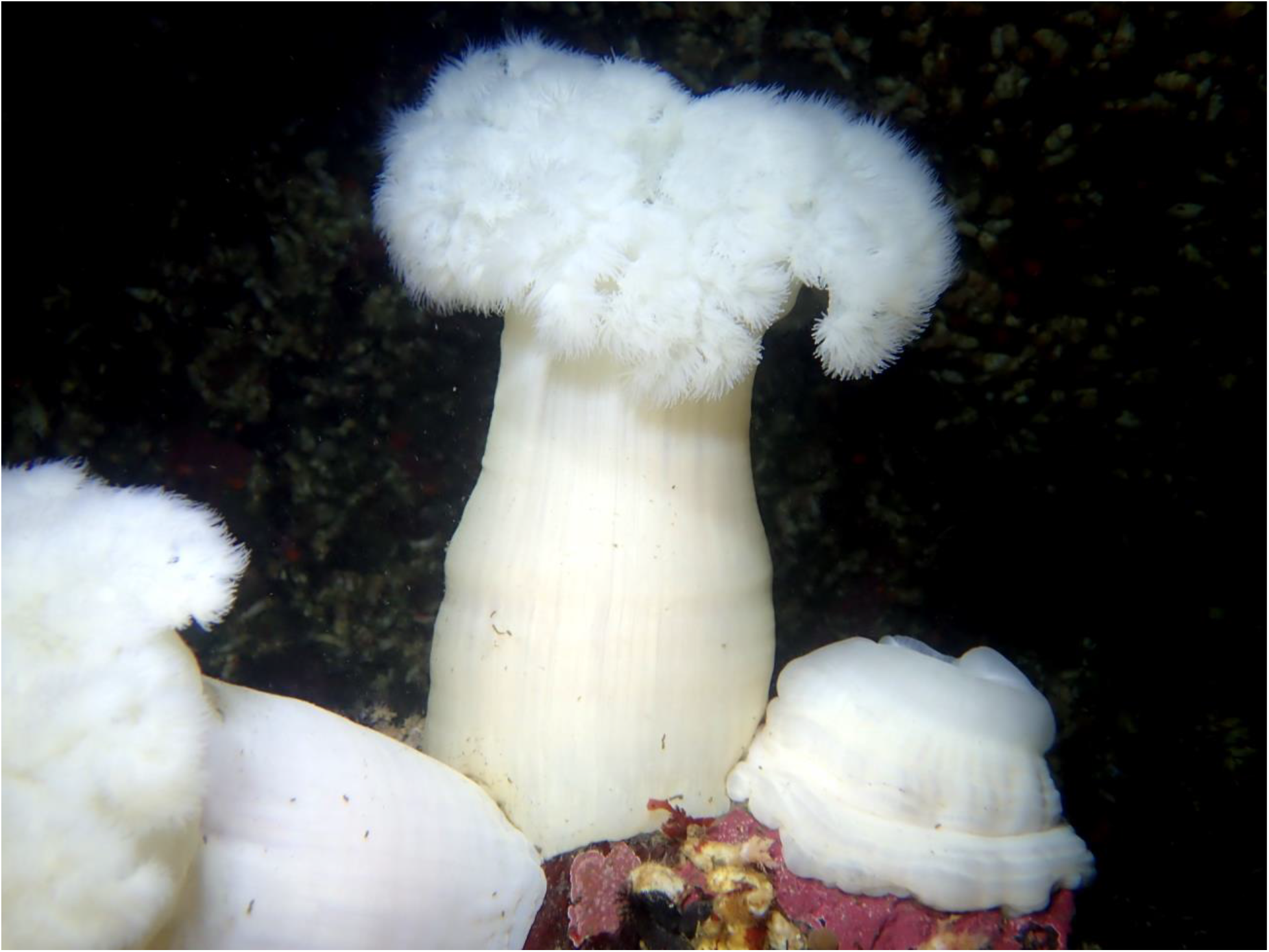
The giant plumose anemone *Metridium farcimen.*

## MATERIALS AND METHODS

### Sample collection and DNA isolation

Plankton communities were quantified by sampling the plankton at three sites within the San Juan Archipelago: adjacent the floating docks of the Friday Harbor Laboratories’ (FHL; Friday Harbor, WA, USA, 48.5452°N, 123.0124°W); 280 m southeast of the docks (48.5436°N, 123.0100°W); and in the San Juan Channel (48.5490°N, 122.9924°W), 1.5 km northeast of the docks. The docks are in an embayment, while San Juan Channel experiences intense tidal flushing and mixing. Samples were taken during an ebb tide between 1:00 and 2:30 pm on August 4, 2016, a season when plankton is highly diverse. Two simultaneous plankton tows were performed at each site at 1-m depth. Each pair of tows included an 80-μm mesh size net to capture a broad range of plankton and a 330-μm mesh size net to capture large zooplankters. Both plankton nets were 50 cm wide, and approximately 98 m^3^ (125 m long tow) of water was sampled. Samples were immediately preserved in 50-mL falcon tubes in 95% ethanol in the field and kept at −20 °C in the laboratory. Falcon tubes were centrifuged at 2000 x g for eight minutes at room temperature to pelletize planktonic particles and remove ethanol. The pellet was homogenized with a mortar and pestle, and the whole tissue homogenate was used for DNA extraction using the MoBIO PowerMax® DNA Isolation Kit following manufacturer’s instructions.

Sixteen *M. farcimen* were collected from the floating docks at FHL by hand one hour after the plankton tows. All collected anemones were within 20 meters of one another at <1 m depth. Anemones were kept in seawater on ice until gut contents could be extracted (between 0.5 and 3 hr following collection). Material attached to the aboral end of the anemones was carefully removed and discarded. In the laboratory, anemones were bisected allowing efficient extraction of gut contents. Material extracted from the gut consisted of partially digested food, copious amounts of mucus, mesenteries, acontia (a structure for agonistic behaviors), and some gonadal tissue in sexually mature individuals. Large food particles (e.g., hydromedusae) were cut up into small pieces to facilitate later grinding. Gut contents were rinsed with 95% ethanol in a 45-μm mesh net to remove excess mucus. Ethanol rinses dissolved anemone mucus more efficiently than seawater. Material within the 45-μm mesh net was further massaged to break up large pieces. During this process, there is a risk of losing partially digested items that lack exoskeletons. The resulting samples were transferred into sterile sample tubes with 95% ethanol and kept at −20 °C overnight. As there were still large particles within the sample, samples were centrifuged at 2000 x g for eight minutes at 20 °C, the supernatant was decanted, and the pellet was ground up within a mortar and pestle into a fine paste. The paste was placed back into another sterile tube with 95% ethanol, centrifuged at 2000 x g for eight minutes at 20 °C, and the supernatant was decanted. The whole pellet of each sample was used for DNA extraction using the MoBIO PowerSoil® DNA Isolation Kit following manufacturer’s instructions. Genomic DNA for both plankton and anemone samples were quantified with a Qubit fluorimeter and diluted to 10 ng/μL.

### PCR and Library Preparation

We used a hierarchical tagging approach with a combination of randomly-assigned tailed PCR primers and single indexed Illumina Y-adapters to sequence all samples in a single Illumina MiSeq run. Three PCR replications were performed per sample. DNA amplification was confirmed on 1.5% gel electrophoresis and then triplicates were pooled. DNA was purified using Solid Phase Reversible Immobilization beads to remove primers, primer dimers, salts and deoxynucleoside triphosphates (dNTPs). DNA samples from the 16 anemones and six plankton samples were used to amplify a highly variable fragment (~313 bp) of the mitochondrial Cytochrome c oxidase subunit I (COI) region with the PCR primers mlCOIintF and jgHCO2198 (Geller, Meyer, Parker, & Hawk, 2013; Leray, Yang, et al., 2013). Despite some amplification bias, this set of primers generates useful estimates of relative abundance (Leray & Knowlton, 2015). PCR reactions were performed in a total volume of 20.0 μL, containing 13.2 μL of nuclease free water, 2.0 μL of Clontech 10X Advantage 2 PCR buffer (Takara Bio Inc., Kusatsu, JP), 1.0 μL of mlCOIintF (10 μM), 1.0 μL of jgHCO2198 (10 μM), 1.4 μL of dNTP, 0.4 μL of Clontech 50X Advantage 2 (Takara Bio Inc., Kusatsu, JP), and 1.0 μL (10 ng) of DNA. The reactions were incubated in a Biometra T3 thermocycler (Analytik Jena, Jena, DE), starting with 5 min of denaturation at 95 °C, followed by 35 cycles of 30 s at 95 °C, 30 s at 48 °C, and 45 s at 72 °C, with a final extension of 72 °C. A negative PCR control and extraction control were performed to test whether the reagents were free of contaminants; both were negative for contamination. Purified PCR products were quantified using an Invitrogen Qubit™ fluorimeter and then diluted to 30 ng/μL. The PCR products of samples amplified with different tailed primers were pooled before library prep as detailed by Leray, Agudelo, Mills, and Meyer (2013) and Leray, Haenel, and Bourlat (2016). Samples were prepared for sequencing with the Illumina TruSeq DNA PCR-free LT Library Prep Kit, which includes end-repair and dA-tailing chemistry, and then ligated with adapters.

### Bioinformatics

Analysis of the sequence data followed the exact same protocol described in Nguyen et al. (2020). Sequences were demultiplexed and Illumina adapters were trimmed using Flexbar (Roehr, Dieterich, & Reinert, 2017). DADA2 (Callahan, McMurdie, & Holmes, 2017; Callahan et al., 2016) was then used to remove primers, discard low quality sequences and infer exact Amplicon Sequence Variants (ASVs) using the following parameters: maxN = 0, maxEE = c (2, 2), truncQ = 10, trimLeft = 26. DADA2 uses sequence quality scores and abundance information to generate an error model that best fit the data, and subsequently uses the error model to infer ASVs. ASVs, which can differ by as little as one nucleotide, were clustered into operational taxonomic units (OTUs) at a 97% identity threshold using VSEARCH (Rognes, Flouri, Nichols, Quince, & Mahé, 2016). To further improve estimates of alpha and beta diversity, spurious OTUs were removed using the LULU algorithm (Frøslev et al., 2017) (parameters: minimum ratio type = “min”, minimum ratio = 1, minimum match = 84, minimum relative co-occurrence = 0.95). This tool, which uses sequence similarity and co-occurrence patterns, was shown to reduce taxonomic redundancy and improve similarity with the true taxonomic composition of test samples (Frøslev et al., 2017).

Taxonomic names were assigned to each remaining OTU using an iterative approach. First, BLASTn searches were used to compare one representative sequence of each OTU to a database of northeast Pacific DNA barcodes. Many of these species were collected off the floats near the source of *M. farcimen*. An OTU was considered to match a local barcode when the level of sequence similarity was higher than 98%. Second, unidentified OTUs were assigned taxonomic information using the Bayesian Least Common Ancestor Taxonomic Classification method (BLCA, Gao, Lin, Revanna, & Dong, 2017) against a curated database of metazoan mitochondrial gene sequences (Midori-Unique v20180221 available at www.reference-midori.info, Machida, Leray, Ho, & Knowlton, 2017). Assignments with less than 50% confidence were not taken into account. Third, the numerous OTUs that remained unidentified using BLCA were compared to the whole NCBI NT database (May 2018) using BLAST searches (word size = 7; max e-value = 5e-13) and assigned the taxonomy of the lowest common ancestor of the first 100 hits.

### Statistical Analyses

Samples with over 95% *M. farcimen* sequences and less than 7000 amplified sequences were dropped from the analysis. It was assumed that these individuals were either not feeding at the time of collection or prey sequences were hidden by the abundant co-amplification of *M. farcimen* sequences leading to insufficient data to categorize diet. Remaining gut content samples (12 of 16) and the plankton samples were rarified to the lowest number of sequences (7268 sequences). An unequal number of sequences can affect estimates of diversity due to the positive relationship between sample size and number of OTUs. This rarefied dataset was used for all further analyses. All data were analyzed in R version 3.5.2 with the vegan 2.5-4 package (Oksanen et al., 2019; R Core Team, 2018).

To illustrate the sequencing effort, rarefaction curves were built for each for each treatment (i.e., gut contents and the two plankton sizes). A plateauing curve indicates an exhaustive sampling effort.

In this study, richness was defined as the number of different OTUs or taxa within a sample or treatment. Abundance was defined as the number of sequences within a sample or treatment. Evenness was defined as the similarity of frequencies of the abundances of OTUs within a treatment. The incidence of an OTU was defined as the fraction of samples or treatments containing that OTU. Mean evenness for each treatment was calculated using Pielou’s evenness index (Pielou, 1966).

Matrices of community dissimilarity based on the Bray-Curtis index were created using both number of sequences and presence/absence of OTUs (i.e., the Sørensen index). Differences between diet composition of *M. farcimen* (n = 12) and 80-μm (n = 3) and 330-μm (n = 3) filtered plankton communities were tested using permutational multivariate analyses of variance (PERMANOVA, Anderson, 2001) with 9999 permutations. Patterns of species composition were visualized in two-dimensional space using non-metric multidimensional scaling ordination plots (nMDS) with 9999 permutations. Similarity percentage analysis (SIMPER) were used to determine what OTUs were significantly contributing to the Bray-Curtis dissimilarities calculated between groups of samples (Clarke, 1993).

## RESULTS

A total of 3,109,361 high-quality metazoan sequences passed the quality controls with an average of 101,000 sequences per plankton sample and an average of 12,500 non-*Metridium* sequences for *M. farcimen* gut content samples. Four *M. farcimen* gut content samples did not provide a sufficient number of sequences (<7000) and were dropped from further analyses, leaving 12 samples. All further results will be reported using the rarefied dataset. The rarefied dataset had 438 OTUs. Among these, 126 OTUs (29% of all OTUs) were identified to species, 381 OTUs (87% of all OTUs) were identified to at least the phylum level, and 57 OTUs (13% of all OTUs) were unidentified metazoans. 174 OTUs (40% of total OTUs) were identified using the Pacific northeast barcode database with 137 of the OTUs being identified to species. Rarefaction curves for both plankton and *M. farcimen* plateaued, which indicates that a sufficient number of sequences was obtained to estimate richness and composition of each sample (Fig. 2).

**Fig. 2.**
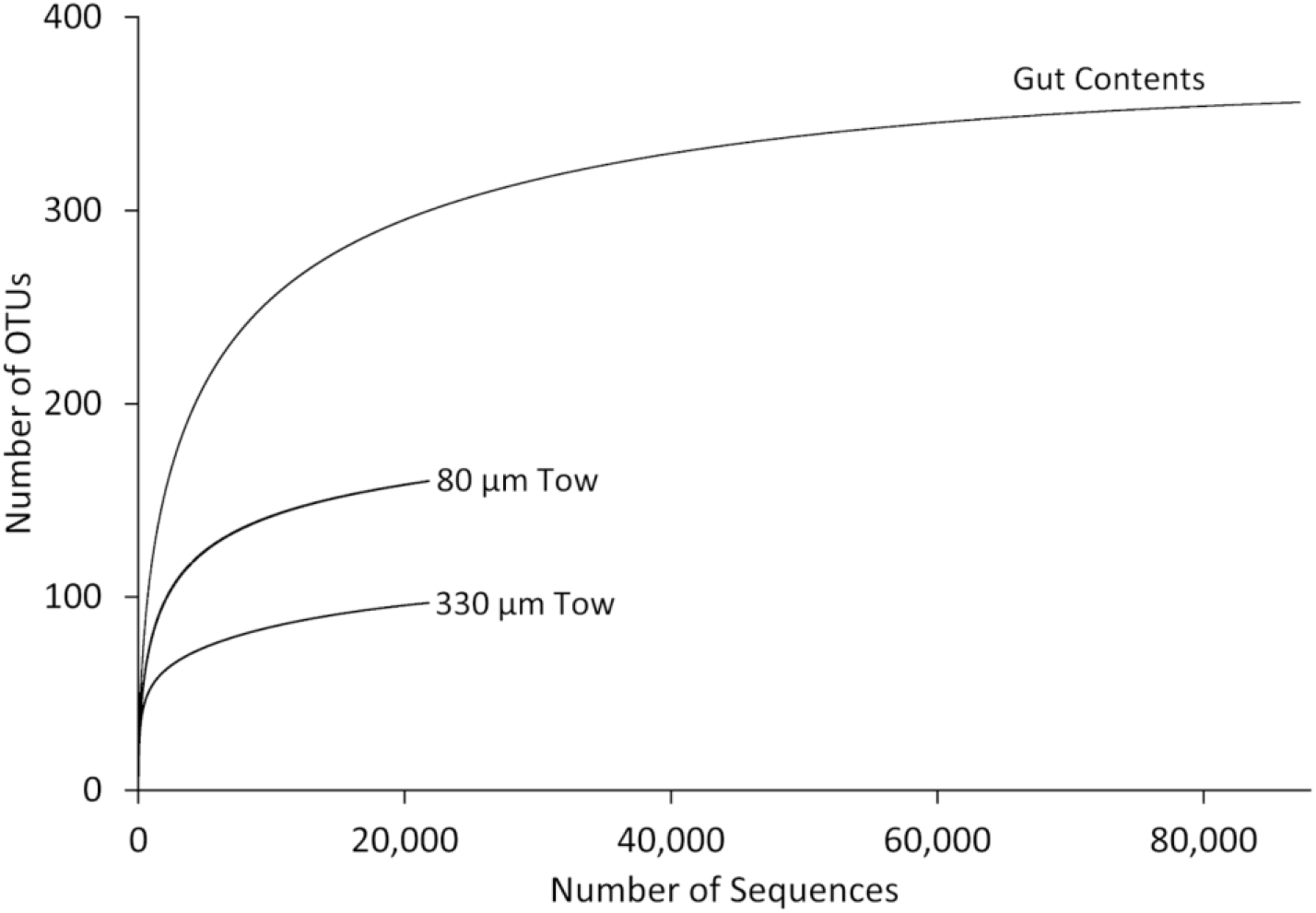
Rarefaction curves to evaluate the completeness of the sequencing effort at describing the diversity of dietary items in the gut contents of *Metridium farcimen* and in nearby plankton samples using two different mesh sizes. All curves plateaued which indicates that this sampling was sufficient.

*M. farcimen* gut contents were richer in total (356 OTUs) than either the 80-μm filtered plankton (160 OTUs) or the 330-μm filtered plankton (97 OTUs). Individual gut contents were less rich on average (74 OTUs) than individual 80-μm filtered plankton samples (91 OTUs) and richer than the individual 330-μm filtered plankton samples (53 OTUs). The 80 and 330-μm filtered plankton samples were more even (0.64 and 0.68, respectively) than the gut contents (0.57). Mean incidence was lowest in the gut contents (21%) and higher in the 80 and 330-μm filtered plankton (57% and 54%, respectively).

The gut contents of *M. farcimen* contained on average 15 classes belonging to 10 animal phyla (Supp. Table 1). *Metridium farcimen* diet contents were primarily made up of arthropods (52% of sequences), especially crabs (presumably larvae) hat this sampling was sufficient, barnacles (larvae or molts), copepods, and insects (Supp. Table 2). Hexanauplia was the most diverse class for both the gut contents and the 80-μm and 330-μm filtered plankton samples (15, 16, and 9 OTUs, respectively) and had the highest proportion of sequences for all three sample types (22%, 38%, and 33%, respectively, Supp. Table 1). The class with the highest incidence within the gut contents of anemones was Hexanauplia (Supp. Table 3). Overall, metabarcoding of gut contents detected many more taxonomic groups than were found in previous conventional visual dietary analysis of *Metridium* spp. conducted by Purcell (1977), Sebens (1981), or Sebens and Koehl (1984). On average, eight metazoan classes were identified in visual identification methods (Purcell, 1977; Sebens, 1981; Sebens & Koehl, 1984) whereas 27 classes were found using metabarcoding (Table 1).

**Table 1.** Comparison of diet composition for *Metridium* species from previous studies and this effort.

| Phylum | Class | Purcell 1977<br>– *M. senile* | Purcell 1977<br>– *M. farcimen* | Sebens 1981<br>– *M. farcimen* | Sebens and<br>Koehl 1984<br>– *M. senile* | This Study<br>– *M. farcimen* |
| --- | --- | --- | --- | --- | --- | --- |
| Annelida | Polychaeta | 15% | 15% | 1% |  | 14% |
| Arthropoda | Arachnida |  |  |  | Present | Present |
|  | Branchiopoda |  |  |  | Present | 4% |
|  | Hexanauplia | 45% | 57% | 87% | 14% | 31% |
|  | Insecta |  |  |  |  | 14% |
|  | Malacostraca | Present | Present | 4% | 65% | 11% |
|  | Ostracoda | Present |  |  | 1% | 7% |
| Bryozoa | Gymnolaemata | Present | 1% | 1% | Present | 3% |
| Chaetognatha | Sagittoidea |  |  |  |  | Present |
| Chordata | Actinopterygii |  |  |  |  | Present |
|  | Ascidiacea |  |  |  | 6% | Present |
| Cnidaria | Anthozoa |  |  |  |  | Present |
|  | Hydrozoa |  |  |  | 12% | 2% |
|  | Scyphozoa |  |  |  |  | Present |
|  | Staurozoa |  |  |  |  | Present |
| Ctenophora | Tentaculata |  |  |  |  | 1% |
| Echinodermata | Asteroidea |  |  | 1% |  |  |
|  | Echinoidea |  |  |  |  | Present |
|  | Holothuroidea |  |  |  |  | 1% |
|  | Ophiuroidea |  |  |  |  | Present |
| Mollusca | Bivalvia | 34% | 30% | 1% |  | 2% |
|  | Gastropoda | 6% | 2% | 1% |  | 2% |
|  | Polyplacophora |  |  |  |  | Present |
| Nematoda | Chromadorea | Present | Present |  | Present | Present |
| Nemertea | Hoplonemertea |  |  |  |  | Present |
|  | Pilidiophora |  |  |  |  | Present |
| Platyhelminthes | Rhabditophora |  |  |  |  | Present |
| Porifera | Demospongiae |  |  |  | Present | 4% |
| Rotifera | Monogononta |  |  |  |  | 3% |

Note: Non-metazoan and unidentified categories (e.g., eggs and embryos) were excluded. Purcell (1977) and Sebens (1981) used relative abundance, Sebens and Koehl (1984) used relative biomass, and we used relative number of sequences. Values less than 0.5% are denoted as “Present”.

There was a significant difference between the communities in the 330-μm plankton samples and the gut samples for both number of sequences (PERMANOVA, *F*_2, 15_ = 1.93, R^2^ = 0.20,*p* < 0.001) and presence/absence (PERMANOVA, *F*_2, 15_ = 2.44, R^2^ = 0.25,*p* < 0.001, Fig. 3-5). There were significant differences between the 80-μm filtered plankton sample and the gut contents for both number of sequences (PERMANOVA, *F*_1, 13_ = 1.31, R^2^ = 0.11, *p* = 0.011) and presence/absence (PERMANOVA, *F*_2, 15_ = 1.72, R^2^ = 0.12, *p* = 0.027). Nineteen and 14 OTUs contributed significantly to the difference between gut content and plankton samples (80-μm and 330-μm, respectively) (SIMPER, p < 0.05, Supp. Table 4-5). All OTUs that differed had higher relative abundances in the plankton than in the gut contents, none had lower abundances. The corrugated clam *Humilaria kennerleyi* (Reeve, 1863), the hydrozoan *Clytia hemisphaerica* (Linnaeus, 1767), the brittle star *Ophiopholis kennerlyi* Lyman, 1860, and the peanut worm *Phascolosoma agassizii* Keferstein, 1866 were all more than 25 times less abundant, relative to other OTUs, in the gut contents compared to the 80-μm filtered plankton. The speckled sanddab *Citharichthys stigmaeus* Jordan & Gilbert, 1882, the periwinkle *Littorina scutulata* Gould, 1849, the bryozoan *Membranipora membranacea* (Linnaeus, 1767), and the hydrozoan *C. hemisphaerica* were all over 70 times less abundant, relative to other OTUs, in the gut contents compared to the 330-μm filtered plankton.

**Fig. 3.**
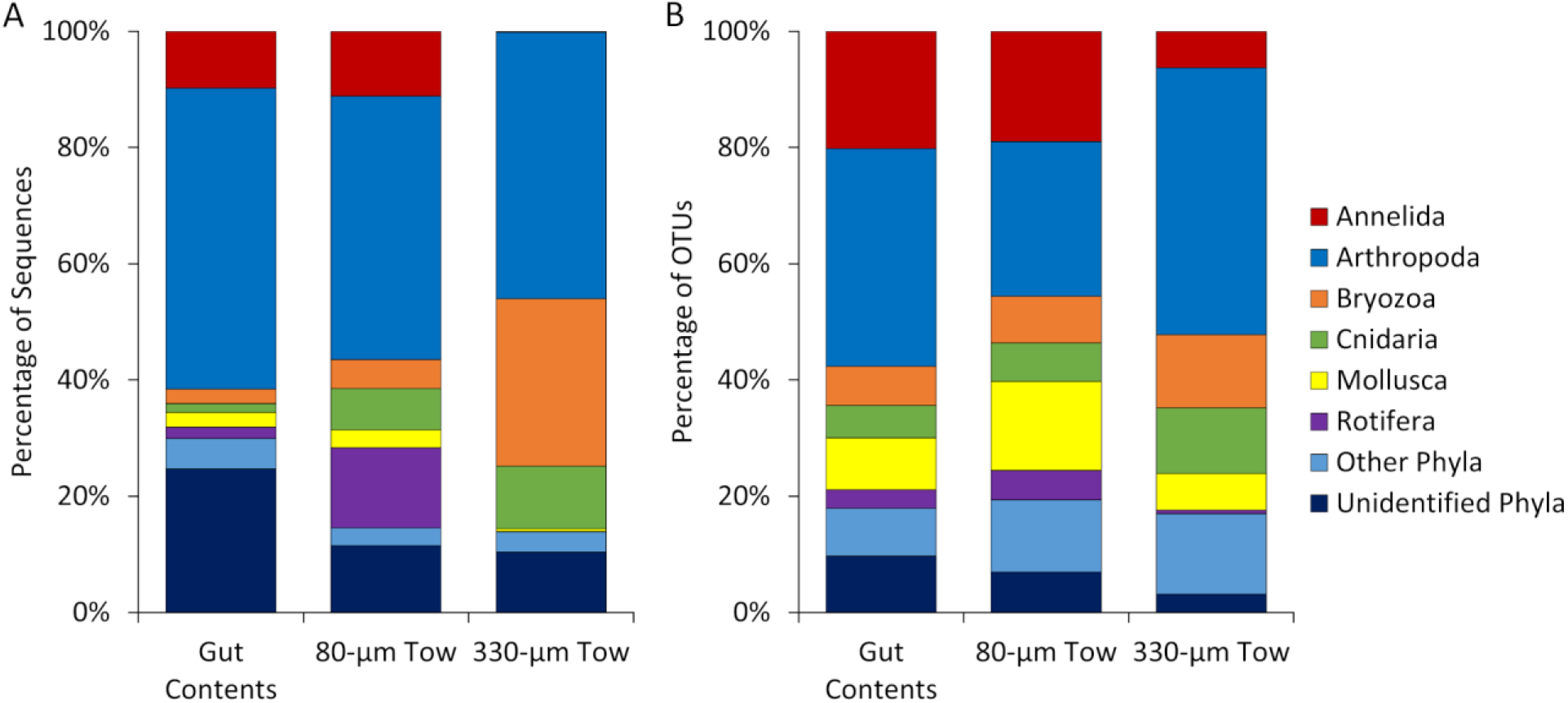
Percentage of (A) sequences and (B) OTUs of major phyla in *Metridium farcimen* gut contents and nearby 80 and 330-μm filtered plankton samples. There were significant differences between the relative abundance of sequences of each community and presence/absence of OTUs (PERMANOVA, p < 0.001).

**Fig. 4.**
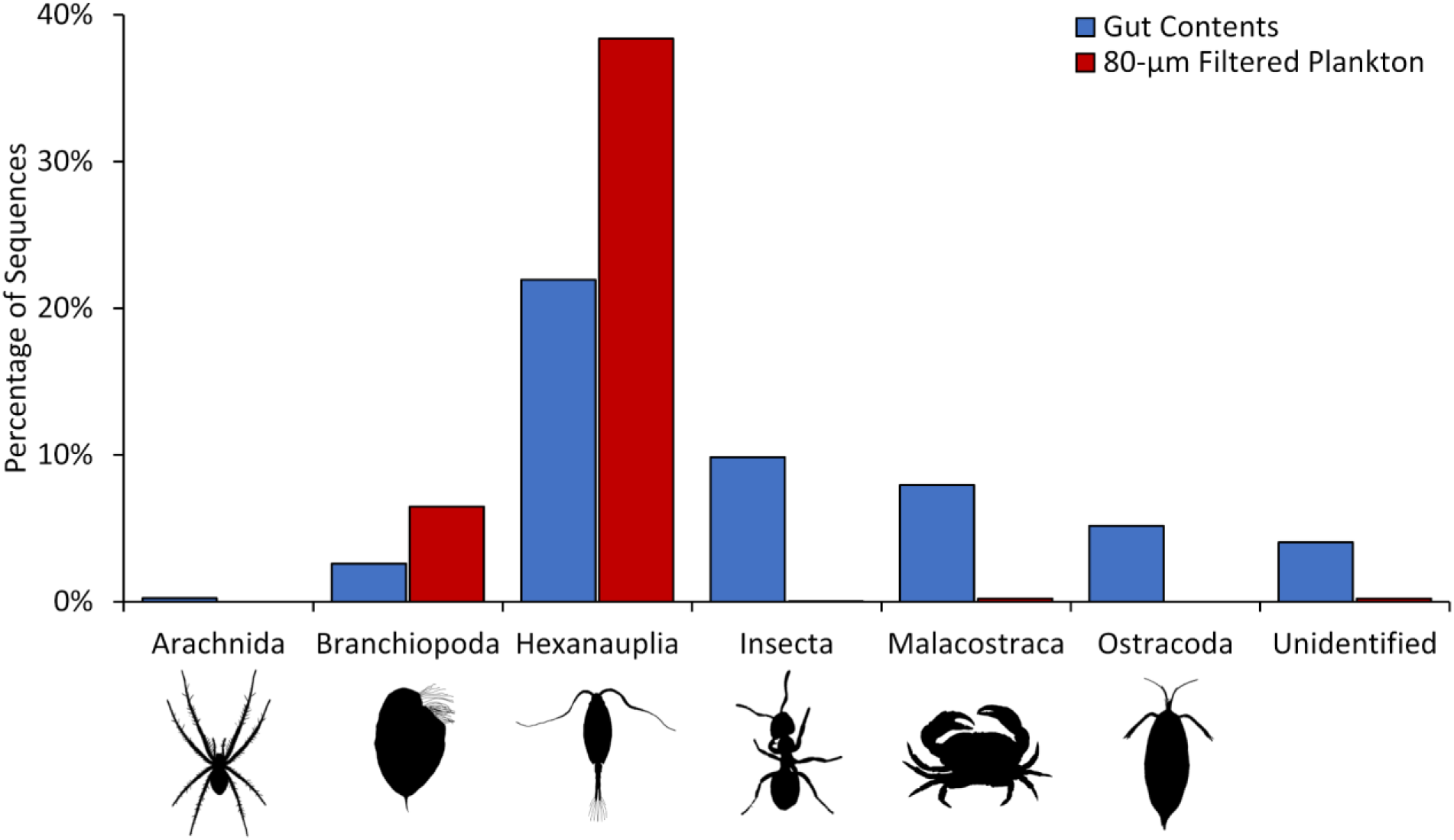
Percentage of sequences of the arthropod classes within the phylum Arthropoda in *Metridium farcimen* gut contents and nearby 80-μm filtered plankton samples.

**Fig. 5.**
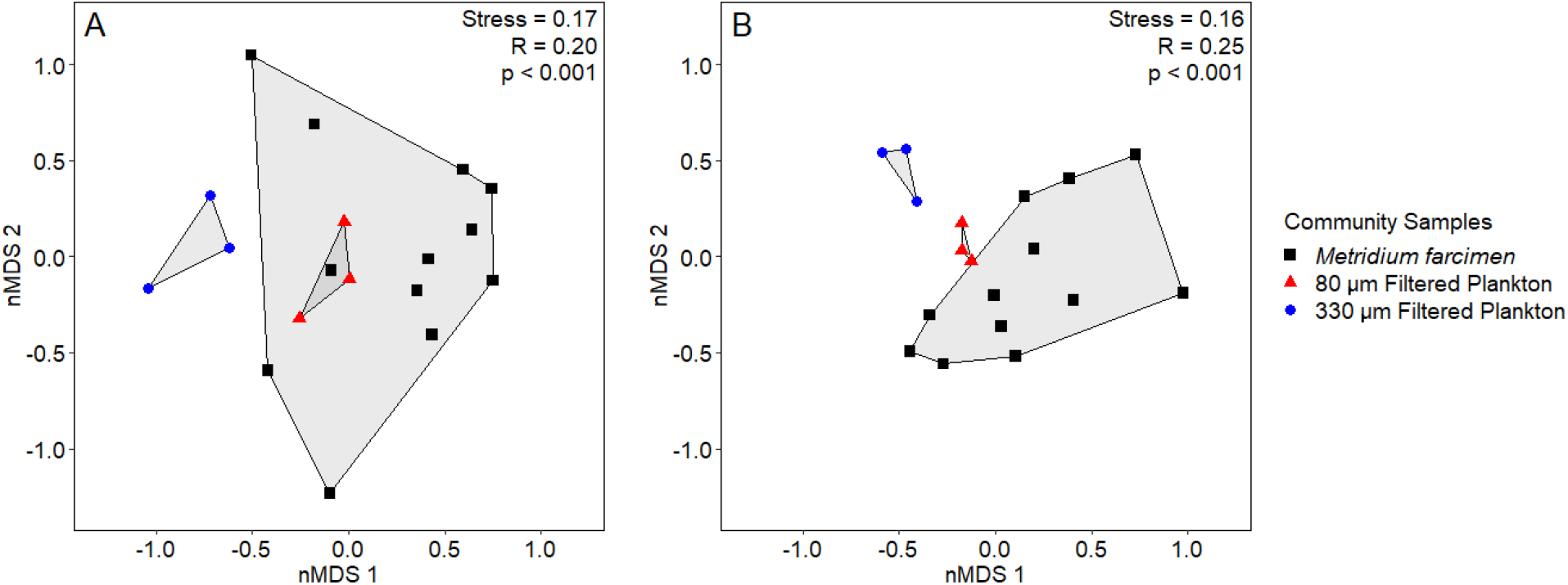
Non-metric multidimensional scaling ordination (nMDS) of (A) number of sequences and (B) presence/absence of OTUs for gut contents of *Metridium farcimen* and nearby plankton samples. Each point represents one sample (plankton tow or gut content). Permutational analysis of variance (PERMANOVA) significance levels are indicated. There were significant differences between the diet of *M. farcimen* and nearby plankton samples.

## DISCUSSION

Overall, *M. farcimen* gut contents held comparable richness to plankton tows on a per-sample basis (mean 71 vs 94/53 OTUs for 80/330 um tows), but held more total diversity (Fig. 2) indicating substantial greater intersample variability. This indicates highly variability in feeding among anemones within 20 m of one another, highlighting the small-scale spatial and temporal heterogeneity in zooplankton availability. A likely reason for this is that in addition to feeding on plankton, *M. farcimen* also captures many benthic taxa that wash off the docks.

It is well known that zooplankton are temporally variable, even at short timescales (Marques, Pardal, Mendes, & Azeiteiro, 2011; Rodríguez et al., 2000). For example, to exclude the temporal variation component while additionally avoiding the issue of variable rates of digestion, Hansson (2006) released and recaptured lab-starved scyphomedusae and then examined what they had consumed over 20 minutes. This method could be adapted to work with anemones and other benthic suspension feeders by starving them in a lab, and, in the case of anemones, deploying them on panels in the field.

Results suggest diet selectivity in *M. farcimen* as was also found by Sebens and Koehl (1984), but contrary to what was found by Purcell (1977). Sebens and Koehl (1984) found that *Metridium senile* (Linnaeus, 1761) preferentially consumed barnacle cyprids, ascidian larvae, and amphipods, and avoided eggs, copepods, and ostracods, compared to availability. In our study, none of the prey species were significantly more common in the diet of *M. farcimen*, but several were less common (Supp. Table 4-5), suggesting that it was unable to detect or capture them. Dietary selectivity in *M. farcimen* seems to be more dependent on prey escape capabilities than predator preferences; this has also been observed in other anthozoans (Sebens, Vandersall, Savina, & Graham, 1996). This result may seem surprising given the larger total richness in the diet of anemones, but individual gut content richness was lower than the 80-μm filtered plankton and mean incidence was half of both plankton samples. While it is unknown how some potential prey species avoid predation, Heidelberg, Sebens, and Purcell (1997) showed that some, but not all, zooplankters can detect passive suspension feeders in moving water and subsequently avoid predation. In addition, they showed that very small prey (e.g., nauplii) were less susceptible to predation. This may be because nematocyst discharge is affected by both chemical cues and mechanical stimulation (Thorington & Hessinger, 1988; Watson & Hessinger, 1988). Larger prey is more likely to impact tentacles at a higher force, increasing capture probability through mechanical or surface chemical detection and consequently eliciting a nematocyst response.

The diet of *M. farcimen* was compositionally different from the diets found by both Purcell (1977) and Sebens (1981). These discrepancies may be partially explained by spatiotemporal differences among these studies locations. Purcell (1977) worked in Monterey, CA on *M. farcimen* associated with pilings at 8 m depth, while Sebens (1981) worked in Harper, WA, farther into the same sound as this study, but on subtidal pilings at 3 m below the surface. Unlike previous morphological dietary analysis (Purcell, 1977; Sebens, 1981), our results show a high relative abundance of demosponge and rotifers. The discrepancy is likely caused by the difficulty in identifying these groups with conventional visual techniques.

While Hexanauplia was the most diverse, rich, and abundant groups in the gut contents, likely because of its high diversity, richness, and abundance in the plankton (Supp. Table 1), we also found a high relative abundance of insects and ostracods in the gut contents (Table 1, Fig. 3-4). The most abundant insect prey (98% of insect sequences) was the pale-legged field ant *Lasius pallitarsis* Emery, 1893 (100% similarity match to the reference sequence GenBank Accession JN292076). This ant species has mating flights in August (Nonacs, 1990), the same month this study was conducted. It seems that *M. farcimen*, when associated with floating docks, may be getting a significant portion of their diet from episodic input from the nearby terrestrial environment. Strong tidal currents and mixing, however, could provide this resource to shallow subtidal populations on natural rock surfaces as well. This result highlights the need for further sampling across a broader temporal and spatial scale and across depth to better understand where *M. farcimen* and other benthic suspension feeders are deriving their energy.

Amplicon sequence variants were clustered into OTUs if they were 97% similar. This can lead to genetically diverse species being clustered into more than one OTU. In several cases, among our samples, sequences classified into different OTUs were mapped to the same species. This implies either that these named species are complexes of multiple cryptic species or that they hold >3% intraspecific variation, something this study cannot determine. Species with more than one OTU delineated were the bryozoans *Celloporella hyalina* (Linnaeus, 1767) and *M. membranacea*, the hydrozoan *Obelia longissima* (Pallas, 1766), the Pacific littleneck clam *Leukoma staminea* (Conrad, 1837), the sharp nose crab *Scyra acutifrons* Dana, 1851, the sea gooseberry *Pleurobrachia bachei* A. Agassiz, 1860, and the syllid worm *Syllis elongata* (Johnson, 1901).

DNA metabarcoding is an efficient method for identifying even partially digested gut contents of animals. However, the results can only be ecologically interpreted if sequences can be matched to taxonomic groups. Building libraries of reference DNA barcodes is time consuming but essential. Metabarcoding has its own limitations that have been reviewed before (de Sousa, Silva, & Xavier, 2019; Deagle et al., 2019). For example, it does not indicate the life stage of prey organisms, while traditional techniques can. It is also semi-quantitative, whereas visual methods can provide absolute counts or biomass. Therefore, we recommend pairing traditional techniques to identify major patterns and metabarcoding to identify the microscopic and partially digested prey items for future intensive studies. We also recommend either using starved animals or sampling the available plankton over a longer time period that best correspond with their prey retention time during digestion.

This work provides important insight into the diet of a competitively dominant sea anemone. Metabarcoding showed that these animals captures a wider range of prey than previously suspected based on conventional visual analysis. The surprising terrestrial input into the diet of *M. farcimen* highlights the need to consider land-sea interactions in trophic models.

## ACKNOWLEDGEMENTS

This work was completed with permission from the director of Friday Harbor Laboratories (FHL). We were funded by the Patricia L. Dudley Endowment for FHL (awarded to CDW), the Kenneth P. Sebens Endowed Student Support Fund (awarded to CDW), the Richard and Megumi Strathmann Fellowship (awarded to CDW), the Robert T. Paine Experimental and Field Ecology Award Fellowship (awarded to CDW), and the FHL Marine Science Fund and Research Fellowship Endowment (both awarded to CDW). We thank the FHL director, faculty, and staff for laboratory space and logistical support; C. Alexandra, A. Ames, M. Ferguson, T. Ferreira, M. Hoban, K. Larkin, K.M. Markello, and J.K. Perez for field and laboratory assistance, and E.R. Anderson, M.N. Dethier, D. Grünbaum, Á. Martínez-Quintana, R. McLachlan, J.L. Ruesink, K.P. Sebens, K.J. Tonra, and M. Turner for reading and commenting on the manuscript.

## AUTHOR CONTRIBUTIONS

CDW, GP and ML conceived and designed the study and collected the data. CDW and BNN analyzed the sequence dataset. Visualizations, statistical analysis, and interpretation were done by CDW and ML. CDW wrote the manuscript.

## DATA ARCHIVING

The demultiplexed and adapter trimmed FASTQ files can be accessed on the NCBI SRA under the BioProject accession number PRJNA682207.

