## Supplemental Table 1-5 for "DNA metabarcoding provides insights into the diverse diet of a dominant suspension feeder, the giant plumose anemone *Metridium farcimen*"

Running title: Metabarcoding and the diverse diet of an anemone

Christopher D. Wells [1*], Gustav Paulay [2], Bryan N. Nguyen [3,4] and Matthieu Leray [5]

1. Department of Geology, University at Buffalo, State University of New York, Buffalo, NY 14260-1300, USA, ORCID: 0000-0002-5945-6171
2. Florida Museum of Natural History, University of Florida, Gainesville, Florida 32611-2062, USA ORCID: 0000-0003-4118-9797
3. Computational Biology Institute, The Milken Institute School of Public Health, George Washington University, Washington, D.C. 20037-2301, USA, ORCID: 0000-0001-7791-4693
4. Department of Biological Sciences, George Washington University, Washington, D.C. 20052-0066, USA
5. Smithsonian Tropical Research Institute, Smithsonian Institution, Panama City 0843-03092, Republic of Panama, ORCID: 0000-0002-7327-1878

*

Supp. Table 1. Average proportion of sequences from plankton samples and the gut contents of *Metridium farcimen* with average number of OTUs per sample in parentheses.

| Phylum | Class | *Metridium farcimen* Gut Contents | 80-µm Filtered Plankton | 330-µm Filtered Plankton |
|---|---|---|---|---|
| Annelida | Polychaeta | 10% (14.4) | 11% (16.3) | Present (1) |
| | Unidentified | Present (0.5) | Present (1) | |
| Arthropoda | Arachnida | Present (0.7) | | Present (0.3) |
| | Branchiopoda | 3% (1.7) | 6% (2.7) | 7% (2.7) |
| | Hexanauplia | 22% (14.5) | 38% (15.7) | 33% (9.3) |
| | Insecta | 10% (1.3) | Present (1) | Present (0.7) |
| | Malacostraca | 8% (3.4) | Present (2) | 5% (6.7) |
| | Ostracoda | 5% (0.9) | | Present (0.3) |
| | Unidentified | 4% (5.3) | Present (3) | Present (1.3) |
| Bryozoa | Gymnolaemata | 2% (4.5) | 5% (7.3) | 29% (8) |
| | Unidentified | Present (0.4) | | |
| Chaetognatha | Sagittoidea | Present (0.2) | Present (2.7) | 2% (4.7) |
| Chordata | Actinopterygii | Present (0.3) | Present (0.3) | 1% (2.3) |
| | Ascidiacea | Present (0.1) | | |
| Cnidaria | Anthozoa | Present (0.2) | | |
| | Hydrozoa | 1% (3.1) | 7% (5.3) | 6% (6.7) |
| | Scyphozoa | Present (0.8) | Present (0.7) | 5% (0.7) |
| | Staurozoa | Present (0.1) | | |
| | Unidentified | Present (0.1) | | |
| Ctenophora | Tentaculata | 1% (0.8) | | Present (1.3) |
| Echinodermata | Echinoidea | Present (0.5) | Present (1.7) | Present (0.3) |
| | Holothuroidea | Present (0.5) | Present (0.7) | |
| | Ophiuroidea | Present (0.1) | Present (2.3) | |
| Entoprocta | Unidentified | | | Present (0.3) |
| Mollusca | Bivalvia | 1% (2.8) | 3% (10) | Present (0.3) |
| | Gastropoda | 1% (3.7) | Present (3.7) | Present (2.3) |
| | Polyplacophora | Present (0.2) | Present (0.3) | Present (0.3) |
| Nematoda | Chromadorea | Present (0.5) | | |
| Nemertea | Hoplonemertea | Present (0.3) | Present (0.3) | |
| | Pilidiophora | | Present (0.3) | |
| Phoronida | Unidentified | | Present (0.3) | |
| Platyhelminthes | Rhabditophora | Present (0.6) | Present (0.7) | |
| | Unidentified | Present (0.9) | Present (0.7) | |
| Porifera | Demospongiae | 3% (1.3) | 2% (1) | Present (0.7) |
| | Homoscleromorpha | | Present (0.3) | |
| | Unidentified | Present (0.2) | | |
| Rotifera | Monogononta | 2% (2.3) | 14% (4.7) | Present (0.7) |
| Unidentified Animal | Unidentified | 25% (7.2) | 11% (6.3) | 10% (1.7) |

Note: Proportions of sequences in *M. farcimen* gut contents most closely resembled the 80-µm filtered plankton except with elevated levels of the classes Malacostraca and Ostracoda. Average proportion of sequences denoted as present are less than 0.5% of the total composition.

Supp. Table 2. The 15 most abundant OTUs in the gut contents of the giant plumose anemone *Metridium farcimen* (68% of sequences).

| Lowest Taxonomic Group | Class | Avg. No. of Sequences |
|---|---|---|
| *Lasius pallitarsis* | Insecta | 703 |
| Metazoa sp. 15 | N/A | 562 |
| Metazoa sp. 1 | N/A | 524 |
| *Metacarcinus gracilis* | Malacostraca | 450 |
| *Balanus* sp. 1 | Hexanauplia | 437 |
| Metazoa sp. 2 | N/A | 356 |
| *Pseudocalanus newmani* | Hexanauplia | 322 |
| Spionidae sp. 1 | Polychaeta | 285 |
| Metazoa sp. 3 | N/A | 240 |
| *Euphilomedes producta* | Ostracoda | 216 |
| *Halichondria panicea* | Demospongiae | 193 |
| *Evadne nordmanni* | Branchiopoda | 185 |
| *Balanus nubilus* | Hexanauplia | 174 |
| *Laonice cirrata* | Polychaeta | 171 |
| Arthropoda sp. 2 | N/A | 162 |

Supp. Table 3. OTUs that were present in more than half of the giant plumose anemone *Metridium farcimen* sampled.

| Lowest Taxonomic Group | Class | No. of *Metridium* Containing |
|---|---|---|
| *Fusitriton magellanicus* | Gastropoda | 9/12 |
| Calanoida sp. 1 | Hexanauplia | 9/12 |
| *Obelia longissima* 1 | Hydrozoa | 9/12 |
| Ploima sp. 1 | Monogononta | 9/12 |
| *Balanus glandula* | Hexanauplia | 8/12 |
| *Tisbe* sp. 1 | Hexanauplia | 8/12 |
| Metazoa sp. 2 | NA | 8/12 |
| *Prionospio lighti* | Polychaeta | 8/12 |
| *Macoma lipara* | Bivalvia | 7/12 |
| *Halichondria panicea* | Demospongiae | 7/12 |
| *Amonardia perturbata* | Hexanauplia | 7/12 |
| *Hydractinia* sp. 1 | Hydrozoa | 7/12 |
| *Diopatra ornata* | Polychaeta | 7/12 |
| *Phyllodoce pettiboneae* | Polychaeta | 7/12 |
| Polycladida sp. 1 | Rhabditophora | 7/12 |

Supp. Table 4. OTUs that were significantly less common in gut contents of the giant plumose anemone *Metridium farcimen* than 80-µm filtered plankton samples (SIMPER, $p < 0.05$).

| Lowest Taxonomic Group | Class | Fold Decrease |
|---|---|---|
| Acartiidae sp. 1 | Hexanauplia | 121 |
| Hexanauplia sp. 1 | Hexanauplia | 65 |
| *Phascolosoma agassizii* | Polychaeta | 41 |
| *Podarkeopsis glabrus* | Polychaeta | 36 |
| *Ophiopholis kennerlyi* | Ophiuroidea | 32 |
| *Humilaria kennerleyi* | Bivalvia | 31 |
| *Clytia languida* | Hydrozoa | 28 |
| Ploima sp. 4 | Monogononta | 20 |
| Ploima sp. 2 | Monogononta | 18 |
| Terebellidae sp. 1 | Polychaeta | 12 |
| *Platynereis bicanaliculata* | Polychaeta | 9 |
| *Notomastus* sp. 1 | Polychaeta | 9 |
| *Leukoma staminea* OTU 2 | Bivalvia | 8 |
| *Nuttalia obscurata* | Bivalvia | 8 |
| *Keenocardium blandum* | Bivalvia | 7 |
| Ploima sp. 1 | Monogononta | 7 |
| *Calanus pacificus* | Hexanauplia | 5 |
| Spionidae sp. 1 | Polychaeta | 5 |
| *Pseudocalanus newmani* | Hexanauplia | 2 |

Supp. Table 5. OTUs that were significantly less common in gut contents of the giant plumose anemone *Metridium farcimen* than 330-µm filtered plankton samples (SIMPER, $p < 0.05$).

| Lowest Taxonomic Group | Class | Fold Decrease |
|---|---|---|
| *Clytia hemisphaerica* | Hydrozoa | 139 |
| *Membranipora membranacea* OTU 3 | Gymnolaemata | 124 |
| *Membranipora membranacea* OTU 2 | Gymnolaemata | 114 |
| *Membranipora membranacea* OTU 5 | Gymnolaemata | 100 |
| *Membranipora membranacea* OTU 4 | Gymnolaemata | 98 |
| *Citharichthys stigmaeus* | Actinopterygii | 74 |
| *Littorina scutulata* | Gastropoda | 72 |
| *Acartia californiensis* | Hexanauplia | 10 |
| Calanoida sp. 2 | Hexanauplia | 9 |
| *Obelia dichotoma* | Hydrozoa | 8 |
| *Diosaccus spinatus* | Hexanauplia | 6 |
| Arthropoda sp. 3 | N/A | 5 |
| *Pleurobrachia bachei* OTU 1 | Tentaculata | 5 |
| Calanoida sp. 1 | Hexanauplia | 3 |
